# Identification of residues in Lassa virus glycoprotein 1 involved in receptor switch

**DOI:** 10.1101/2021.04.30.442226

**Authors:** Jiao Guo, Xiaoying Jia, Yang Liu, Junyuan Cao, Gengfu Xiao, Wei Wang

**Author notes:** Address correspondence to Wei Wang.

## Abstract

Lassa virus (LASV) is an enveloped, negative-sense RNA virus that causes Lassa hemorrhagic fever, for which there are limited treatment options. Successful LASV entry requires the viral glycoprotein 1 (GP1) to undergo a receptor switch from its primary receptor alpha-dystroglycan (α-DG) to its endosomal receptor lysosome-associated membrane protein 1 (LAMP1). A conserved histidine triad in LASV GP1 has been reported to be responsible for receptor switch. To test the hypothesis that other non-conserved residues also contribute to receptor switch, we constructed a series of GP1 mutant proteins and tested them for binding to LAMP1. Four residues, L84, K88, L107, and H170, were identified as critical for receptor switch. Substituting any of the four residues with the corresponding lymphocytic choriomeningitis virus residue (L84N, K88E, L10F, and H170S) reduced the binding affinity of GP1_LASV_ for LAMP1. Moreover, all the mutations caused decreases in GPC-mediated membrane fusion at both pH 4.5 and 5.2. The infectivity of pseudotyped viruses bearing either GPC^L84N^ or GPC^K88E^ decreased sharply in multiple cell types, whereas L107F and H170S had only mild effects on infectivity. Notably, in LAMP1 knockout cells, all four mutants showed reduced pseudovirus infectivity. Using biolayer light interferometry assay, we found that all four mutants had decreased binding affinity to LAMP1, in the order L84N > L107F > K88E > H170S.

**IMPORTANCE:** Lassa virus requires pH-dependent receptor switch to infect host cells; however, the underlying molecular mechanisms of this process are not well known. Here, we identify four residues, L84, K88, L107, and H170 that contribute to the interaction with the second receptor lysosome-associated membrane protein 1 (LAMP1). Mutant any of the four residues would impair the binding affinity to LAMP1, decrease the glycoprotein mediated membrane fusion, and reduce the pseudovirus infectivity.

## INTRODUCTION

Lassa virus (LASV) belongs to the *Arenaviridae* family of enveloped, negative-sense, bi-segmented RNA viruses and is classified as an Old World (OW) mammarenavirus (1). It is transmitted from the rodent host *Mastomys natalenis* to humans via contaminated excreta (2). LASV infections cause Lassa hemorrhagic fever, which causes high mortality in hospitalized patients. However, there are no FDA-approved vaccines or specific antiviral agents against LASV (3).

The LASV RNA genome encodes an RNA-dependent RNA polymerase (L), nucleoprotein (NP), matrix protein (Z), and a highly glycosylated membrane glycoprotein (GP). GPC is synthesized as an inactive precursor glycoprotein complex, then cleaved into three subunits: a stable signal peptide (SSP), a receptor-binding subunit (GP1), and a membrane-spanning fusion subunit (GP2) (4). During cellular entry, LASV GP1 interacts with the primary receptor, alpha-dystroglycan (α-DG), on the plasma membrane; LASV is then internalized through macropinocytosis, subsequently reaching the late endosomal compartment (5). In this low pH environment, GP undergoes irreversible structural changes that decrease its affinity for α-DG, while increasing its affinity for the acidic receptor, lysosome-associated membrane protein 1 (LAMP1) (5–8). A histidine triad (H92/93/230) in GP1_LASV_ is highly conserved among OW mammarenaviruses and is indispensable for LAMP1 binding (9). Several other residues in LASV GP1 that are not shared by other OW mammarenaviruses have also been reported to be critical for LAMP1 binding (10). Here, we focused on comparing LASV with the prototypical OW mammarenavirus lymphocytic choriomeningitis virus (LCMV), which is genetically and serologically similar to LASV but does not interact with LAMP1. We mutated several residues in GP1_LASV_ to their corresponding residues in GP1_LCMV_, tested its ability to bind to LAMP1, and found four residues involved in LAMP1 binding.

## RESULTS

### Four single-residue substitutions in GP1_LASV_ disrupt LAMP1 binding

By comparing the amino-acid sequences of GP1_LASV_ and GP1_LCMV_ (Fig. 1A), several residues located at the tip of GPC_LASV_ or spatially close to the histidine triad (H92/93/230) were selected and mutated to the corresponding residue in GP1_LCMV_. LAMP1 binding was evaluated using pull-down assays at pH 5.0. As shown in Fig. 1B, four mutants, L84N, K88E, L107F, and H170S, disrupted the binding of GP1_LASV_-Fc to LAMP1, wherease other mutants, such as T87S, E100S, T101S, L105P, F147N, K161E, K161Q, and D229K, had little effect. H92Y, H93Y, and H230Y were used as the positive controls, none of which showed binding to LAMP1, consistent with the previous reports (9, 11).

**Fig. 1.**
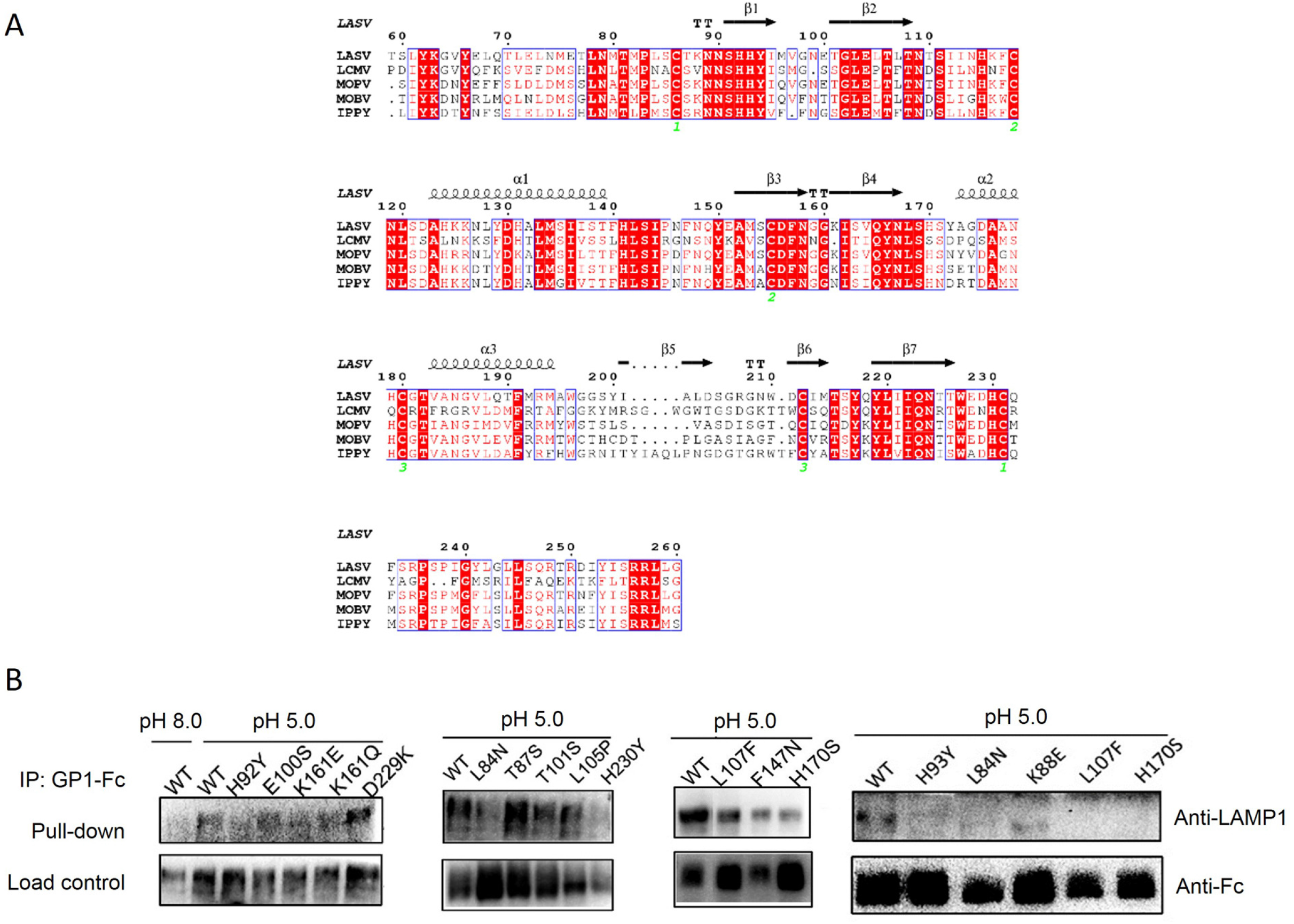
Binding ability of the mutant GP1_LASV_ to LAMP1. **(A)** Multiple sequence alignment of GP1 proteins of representative OW mammarenaviruses. The UniProt accession codes of the sequences are P08669 (LASV), P07399 (LCMV), P19240 (MOPV), Q2A069 (MOBV), and Q27YE4 (IPPY). Fully conserved residues were highlighted with a red background, and partially conserved residues were presented in red. The secondary structure observed with GP1_LASV_ (PDB: 4ZJF) was indicated above the sequence, and cysteine involved in the disulfide bond was numbered below the alignment in green. This graphical representation was generated using ESPript (http://espript.ibcp.fr). **(B)** Images of LAMP1 pull-down assays by WT and mutated GP1_LASV_-Fc. GP1_LASV_-Fc was immobilized on sepharose beads, incubated with equal amount of cell lysates at pH 5.0 and pH 8.0, pulled down, and eluted. Protein pellets were recovered and subjected to western blot analysis.

### Effects of the mutants on GPC cleavage efficiency

To evaluate the effect of the four mutations on the cleavage processing of LASV GPC into GP1 and GP2, 293T cells were transfected with pCAGGS vectors encoding WT and the mutant GPCs. As shown in Fig. 2, GPC^L107F^ and GPC^H170S^ exhibited cleavage efficiencies similar to that of GPC^WT^, whereas GPC^L84N^ and GPC^K88E^ showed cleavage processing defects (Fig. 2A). Compared with GPC^WT^, the cleavage efficiency of GPC^L84N^ decreased to only approximately 4% of that of GPC^WT^. The cleavage efficiency of GPC^K88E^ was approximately 25% that of GPC^WT^ (Fig. 2B). There were no significant differences in the cleavage efficiencies of GPC^L107F^, GPC^H170S^, and GPC^WT^, indicating that unlike L84N and K88E, L107F and H170S might have little effect on the cleavage efficiency of GPC.

**Fig. 2.**
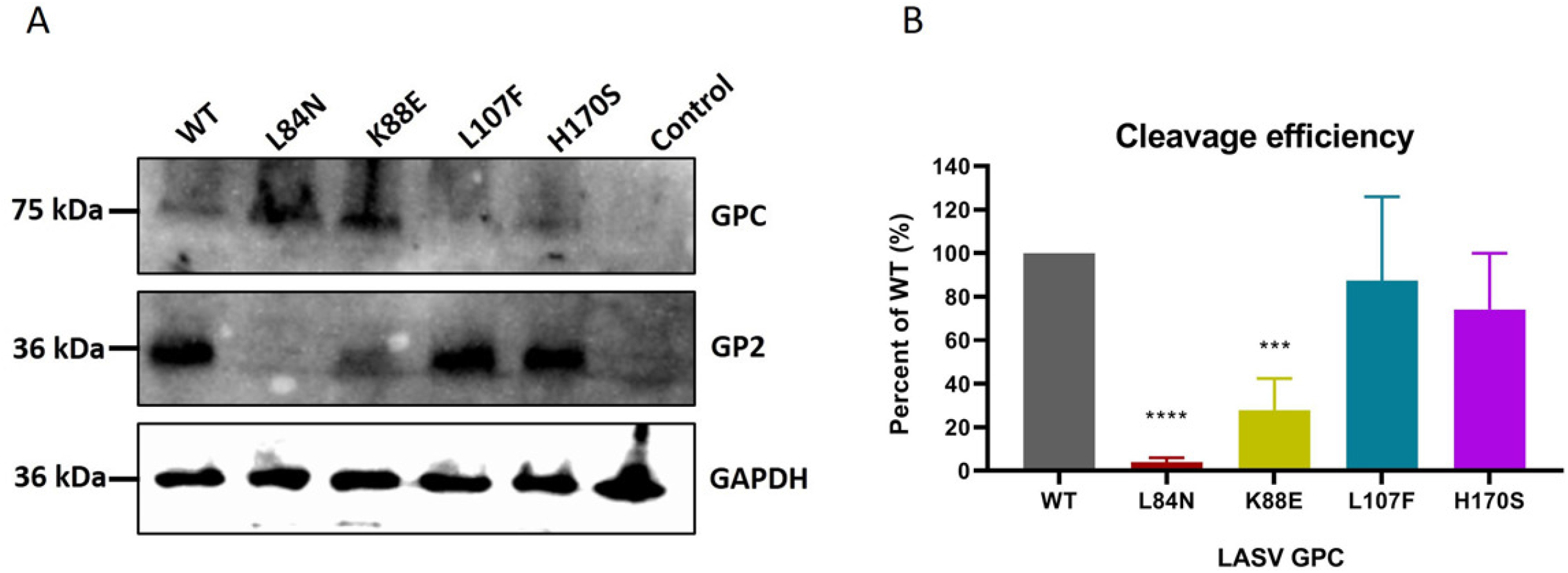
Cleavage processing of WT and mutant LASV GPCs. (A) 293T cells were transfected with WT or one of the four GPC mutants (L84N, K88E, L107F, and H170S).. The equal cell lysates were subjected to western blotting and probed with an anti-LASV GP2 antibody. The image was representative of three independent experiments. (B) Quantitative analysis of the cleavage efficiencies based on the intensity of the bands. Data were presented as means ± SDs from three independent experiments. \*\*\*\**P* < 0.0001, \*\*\**P* < 0.001.

### Effects of mutants on GPC-mediated membrane fusion

GPC cleavage is upstream of GPC-mediated membrane fusion, so a decrease in cleavage efficiency would lead to a reduction in the fusion activity. To this end, the effects of the substitutions on GPC-mediated membrane fusion were also tested. As shown in Fig. 3, the efficiency of membrane fusion mediated by GPC^L84N^ was lower than that of GPC^WT^ (< 20%) at all tested pH values (4.5, 5.0, and 5.2). Compared with GPC^WT^, the membrane fusion efficiency of GPC^K88E^ ranged from 42 to 63%, while that of GPC^L107F^ ranged from 46 to 80%. The decrease in fusogenicity of L84N and K88E might be due to a deficiency in GPC cleavage. Notably, the membrane fusion efficiency of GPC^H170S^ was nearly twice as high as that of GPC^WT^ at pH 5.0. However, the membrane fusion efficiency of GPC^H170S^ was lower than that of GPC^WT^ at both pH 4.5 and pH 5.2, suggesting that pH 5.0 might be an optimal pH for GPC^H170S^-mediated membrane fusion.

**Fig. 3.**
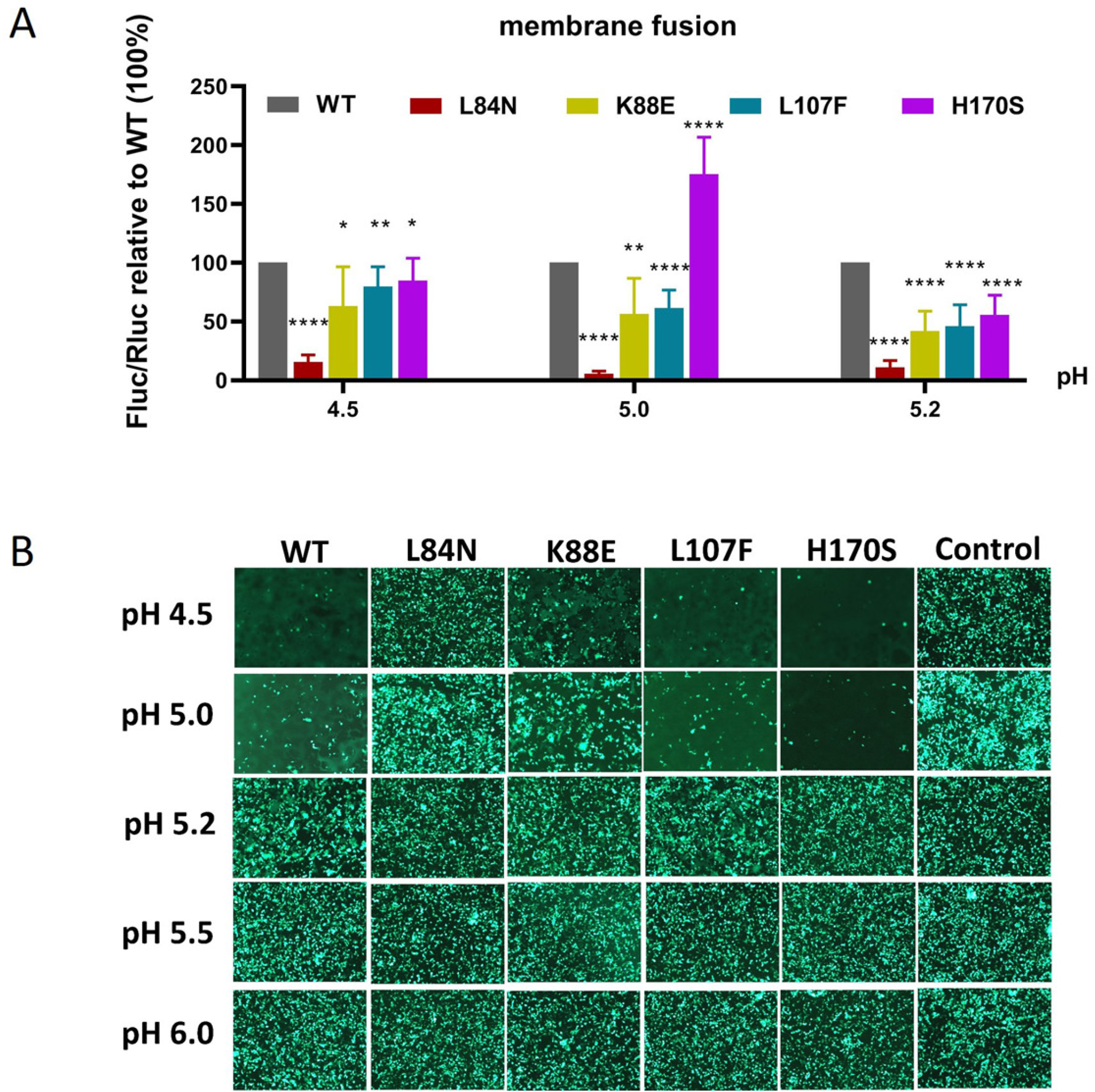
Membrane fusion by WT and four mutant GPCs. (A) Quantification of membrane fusion using a dual-luciferase reporter assay at the indicated pH values. Data are presented as means ± SDs from three independent experiments. \*\*\*\**P* < 0.0001, \*\*\**P* < 0.001, \*\**P* < 0.01, \**P* < 0.05. (B) Qualitative analysis of membrane fusion was performed in 293T cells co-transfected with pEGFP-N1 and pCAGGS-LASV GPC plasmids.

### Four substitutions in GP1_LASV_ reduce LASV infectivity

We constructed LASV pseudotyped viruses (pvs) with a VSV backbone and luciferase reporter gene. The genome copy numbers of LASV_PV_^WT^, LASV_PV_^L84N^, LASV_PV_^K88E^, LASV_PV_^L107F^, and LASV_PV_^H170S^ were 8.10 × 10^9^, 2.05 × 10^9^, 1.90 × 10^10^, 1.18 × 10^10^, and 7.88 × 10^9^ per mL, respectively.

The WT and mutant viruses with the same copy number were used to infect different cell lines, and relative luminescence was measured to evaluate efficiency. In A549 and BHK cells, L84N, K88E, L107F, and H170S significantly reduced infectivity relative to WT, The infection efficiency of L84N was reduced by 3.3 log values. In HEK 293T cells, L84N and K88E significantly reduced infectivity, whereas L107F and H170S had little effect. In Vero cells, the infectivity of L84N, K88E, and H170S was significantly reduced, whereas L107F had little effect. In summary, the infectivity of GPC^L84N^ and GPC^K88E^ mutant pvs was sharply decreased in both tested cell lines, whereas L107F and H170S had milder effects on infectivity (Fig. 4).

**Fig. 4.**
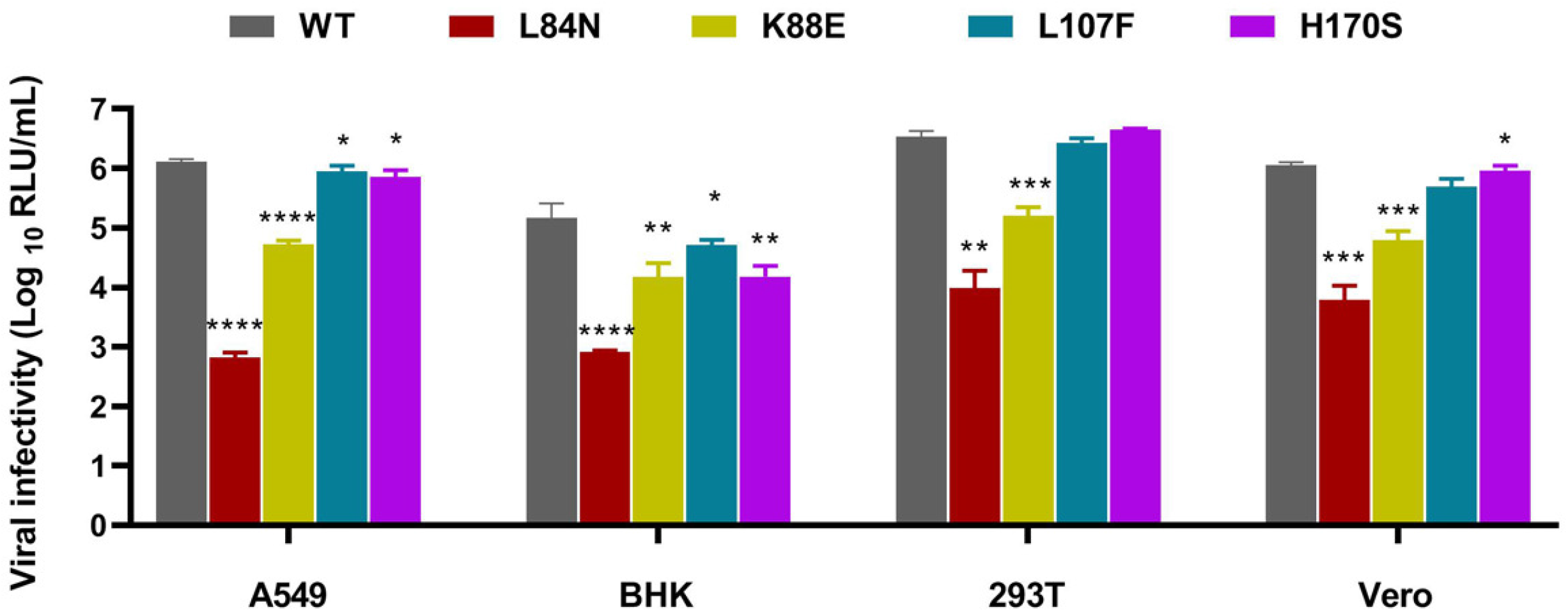
Infectivity of multiple cell lines by the mutant pseudoviruses. Viral copy number was normalized using quantitative RT-PCR. After normalization, A549, BHK, 293T, and Vero cells in 96-wells plate were infected with 2 × 10^6^ copies / well, respectively. After 24 h, the cell lysates were subjected to evaluate the *Renilla* luciferase activities. Data were presented as means ± SDs from three independent experiments. \*\*\*\**P* < 0.0001, \*\*\**P* < 0.001, \*\**P* < 0.01, \**P* < 0.05.

To evaluate the role of LAMP1 in LASV infection, we knocked out A549 cells using CRISPR/Cas9 gene editing, then measured LAMP1 expression was evaluated using western blotting (Fig. 5A). LASV_PV_ replication was reduced in LAMP1 knockout A549 cells compared with that in wild-type A549 cells (Fig. 5B), confirming the indispensable role of LAMP1 in LASV infection. Moreover, compared with LASV_PV_^WT^, all four mutants showed decreased infectivity in LAMP1 KO cells. Intriguingly, the decreases we observed in the infectivity of LASVpv^L107F^ and LASVpv^H170S^ in KO cells were greater than those in WT cells (Fig. 4).

**Fig. 5.**
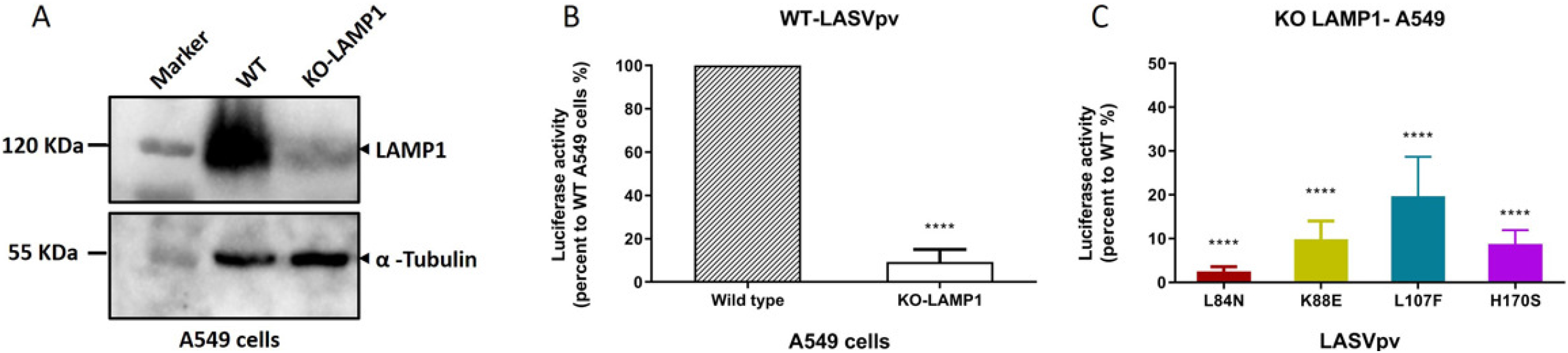
Infectivity of pseudoviruses on WT and LAMP1-KO A549 cells. (A) WT and LAMP1-KO A549 cells were analyzed using western blotting with antibody against LAMP1. (B) Infectivity of LASVpv^WT^ for WT and LAMP1-KO A549 cells. WT and LAMP1-KO A549 cells were infected with LASVpv^WT^ (2 × 10^6^ copies) for 1 h, respectively, the cells were lysed and assayed for RLU at 24 h post-infection. (C) Infectivities of LASVpv^L84N^, LASVpv^K88E^, LASVpv^L107F^, and LASVpv^H170S^ for LAMP1-KO A549 cells. Data are presented as means ± SDs from three independent experiments. \*\*\*\**P* < 0.0001.

**Fig. 6.**
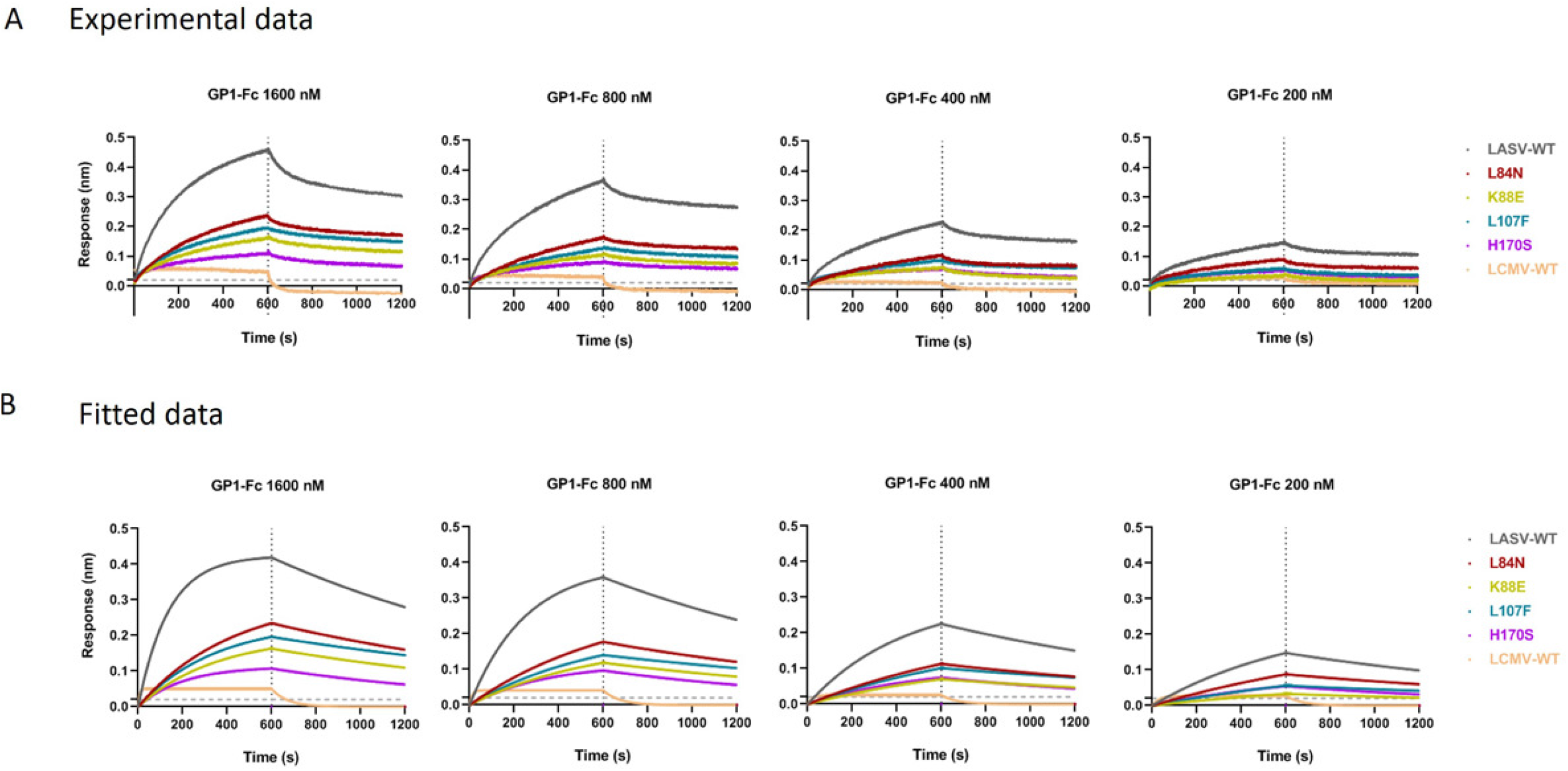
Analysis of GP1-LAMP1 interaction. Binding of WT and mutant GP1_LASV_ to the distal domain of LAMP1 was measured using the Octet RED 96 instrument. Every column presents the maximum values of the binding curves. The response values (nm) to the indicated concentrations of GP1_LASV_ (1600, 800, 400, and 200 nM) were measured. Experimental data are shown in (A); these data were fit with a 1:1 global fitting model and the fitted curves are presented in (B).

### GP1 mutants have reduced binding affinity for LAMP1

To further evaluate the role of the mutated residues (L84N, K88E, L107F, and H170S) in LAMP1 binding, biolayer interferometry (BLI) was performed to test the ability of these mutants to bind to LAMP1 (12). As shown in Fig. 7A, the WT GP1-Fc as well as the entire mutant GP1-Fc exhibited dose-dependent binding to LAMP1, while LCMV GP1-Fc showed no binding. Specifically, the relative LAMP1 binding affinities were L84N > L107F > K88E > H170S (Fig 7A and B). LAMP1 binding was robustly abolished by the H170S substitution; the K88E, L84N, and L107F mutants weakly bound to LAMP1. These results are consistent with those of the LAMP1 pulldown assay, indicating that L84N, K88E, L107F, and H170S have negative effects on LAMP1 binding.

**Fig. 7.**
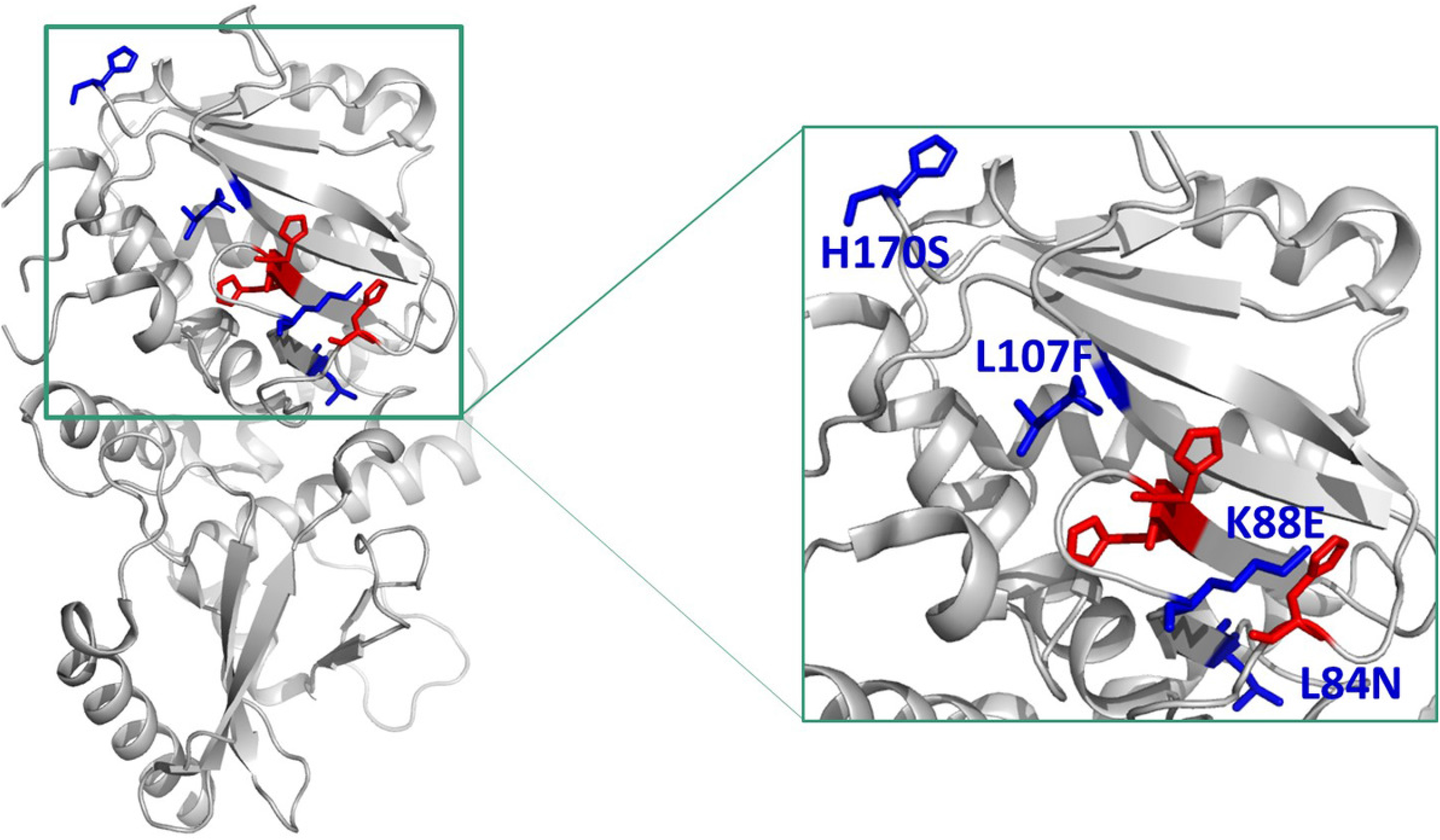
Crystal structure of GP1_LASV_ (PDB: 5VK2), showing the positions L84, K88, L107, H170 (blue) and the histidine triad (red); these residues are magnified in the inset box.

## DISCUSSION

LASV GPC mediates viral entry into host cells; therefore, substantial efforts have been made to understand its structure, function, and immunobiology (13, 14). Successful LASV entry requires a switch in binding from α-DG to the endosomal receptor LAMP1 (8). Ebola virus (EBOV) uses a similar, multistep entry strategy, employing Niemann-Pick type C1 (NPC1) to complete its entry step (15–17). Tetraspanin CD63 was shown to enhance the efficiency of GP-mediated membrane fusion for another OW mammarenavirus Lujo virus (LUJV), and is hypothesized to be a possible second receptor for LUJV entry (18, 19).

The prototypical OW mammarenavirus LCMV is genetically and serologically similar to LASV and also is pathogenic to humans; both share similar receptor-binding domains involved in α-DG binding (20). A previous study implicated a conserved, positively charged histidine triad (H92/93/230) in GP1_LASV_ binding to LAMP1 (9), located on the β-sheet face of GP1_LASV_. GP1_LCMV_ shares this histidine triad, but LCMV does not undergo receptor switch. The GP1 sequence is different between the two, which might lead to differences in receptor binding.

A previous study implicated important fragments in addition to the histidine triad in LASV receptor switch. These residues were mainly located in β5 and L7 of GP1_MORV_ and play key roles in LAMP1 binding. Notably, when partial fragments of GP1_MOPV_ were mutated to their corresponding LASV residues, the resulting chimeric GP1_MORV_ could bind to LAMP1, suggesting that these residues play a key role in the binding of GP1_LASV_ to LAMP1 (10).

Based on the crystal structure of GP1_LASV_ (PDB: 5VK2), K88 is located in the center of the histidine triad (Fig. 7) (4). Notably, the K88E substitution significantly reduced membrane fusion efficiency, suggesting that the charged residue K88 plays an important role in GPC-mediated membrane fusion, which might sense the acidic pH within the endolysosomal compartment, thus promoting receptor switch and the conformational change of GP1_LASV_. Thus, we infer that maximal LAMP1 binding and membrane fusion of LASV require a positive charge at position 88 in GPC_LASV_. The mechanism involving K88 might be similar to that of K33, a key charged residue located in the putative membrane-proximal region of the SSP. It has been reported that charge-changing substitutions of K33 affect the maturation of GPC and its downstream function; thus, the K33 residue is crucial for GPC sensitivity to pH (21). Another positively charged residue, H170, is located on the top of the GP1_LASV_ structure and may function similarly to K88 and K33 in sensing pH. Moreover, L84 and L107 are located in the vicinity of the histidine triad and may be important for maintaining the structure of the histidine triad and the GP1_LASV_-LAMP1 interaction interface.

To investigate whether these substitutions affect the binding of GP1_LCMV_ to LAMP1, we constructed four GP1_LCMV_ mutant expression plasmids (N90L, V94K, F110L, and S174H, according to the residues in GPC_LCMV_). However, none of these GP1_LCMV_ mutants interacted with LAMP1. This suggests that the impact of these four residues in GP1_LASV_ was greater than that in GP1_LCMV_, and that the corresponding mutations in GP1_LCMV_ were not sufficient to promote its interaction with LAMP1.

To date, there is no published structure of LAMP1 cocrystallized with GP1_LASV_; therefore, we performed a BLI assay to evaluate the binding of distal-LAMP1 and GP1_LASV_. Distal-LAMP1 refers to residues 27–194 of LAMP1, the key domain that partially reflects the physiological state and function of LAMP1 in cells; thus, it has been used to study LAMP1 function (22). The data from this assay were consistent with the pulldown data, as distal-LAMP1 plays a critical role in LAMP1 binding.

LAMP1 triggers significant increase in GPC_LASV_ spike complex-mediated cell entry (7). However, we found that LASV infection was not completely inhibited in LAMP1 knockout A549 cells (Fig. 5B). These data are consistent with those from a previous study, in which the infectivity of LASV VSV pseudoviruses was reduced by 70–85% in LAMP1 knockout cells relative to wild-type cells (23).

These four residues, identified as being involved in LASV glycoprotein receptor switch, are significant for understanding the mechanism underlying host-pathogen interactions and provide new insight for further study of invasive mechanisms, and therefore potential inhibitors, of LASV. Further elucidation of the relationship between α-DG and LAMP1 and characterization of the LAMP1-GP1 interaction will clarify the events that occur during LASV infection and facilitate specific strategies to clinically inhibit this crucial interaction (24).

## MATERIALS AND METHODS

### Cell lines, plasmids, and antibodies

293T, BHK, A549, and Vero cells were maintained in Dulbecco’s modified Eagle’s medium (DMEM; Gibco, Grand Island, NY, USA) supplemented with 10% fetal bovine serum (FBS; Gibco). 293F cells were cultured in Freestyle 293 expression medium (Invitrogen, Carlsbad, CA, USA). GPC_LASV_ (Josiah strain, GenBank HQ688673.1) was synthesized by Sangon Biotech (Shanghai, China) and subcloned into the pCAGGS vector. The recombinant pcDNA3.1-GP1_LASV_-Fc plasmid was constructed by inserting codon-optimized GP1_LASV_, which was synthesized de novo by GenScript (Nanjing, China). Distal LAMP1-Fc expressing plasmid was a gift from Professor Ron Diskin (Department of Structural Biology, Weizmann Institute of Science, Rehovot, Israel).

Anti-LAMP1 antibody was purchased from Millipore (Billerica, MA, USA). Anti-GAPDH, anti-α-tubulin, anti-Fc, and horseradish peroxidase (HRP)-conjugated secondary antibodies were obtained from Proteintech (Wuhan, China). Anti-GP2_LASV_ antibodies were produced in our laboratory. (25, 26).

### Protein expression and purification

GP1_LASV_-Fc and distal LAMP1-Fc were transfected into HEK 293F cells using polyethylenimine (PEI) at a density of 2.0 × 10^6^/mL. Overgrowth of the cells was controlled by adding 2 mM sodium valproate and shaking at 120 rpm with 8% CO_2_ at 37°C. After 5 days, suspension cultures were harvested, centrifuged at 3500 × *g* for 15 min, and sterilized by passage through 0.45 μm filters (Thermo Scientific, USA). Secreted proteins were purified from the culture supernatants using a protein A+G affinity column (GE Healthcare) according to the manufacturer’s instructions. Protein concentrations were determined based on absorption at 280 nm (UV_280_) using theoretical extinction coefficients. Protein samples were analyzed using blue native polyacrylamide gel electrophoresis (PAGE) and stained with Coomassie blue to assess purity.

### LAMP1 pulldown assays

Assays were performed as described previously (7, 10). Briefly, LASV GP1-Fc protein was incubated overnight with protein A+G Sepharose beads (ProteinTech) at 4°C and then washed five times with NETI buffer (50 mM Tris-HCl, 1 mM EDTA, 150 mM sodium chloride, 0.5% [vol/vol] IGEPAL, pH 8.0). The beads were incubated with extracts from HEK 293T cells prepared in NETI buffer and adjusted to pH 5.0, at 4°C for 3 h. Proteins were eluted using 20 mM Tris-HCl (pH 8.0) and 150 mM sodium chloride and precipitated using cold acetone at −20°C. Protein pellets were recovered in sample buffer and subjected to western blotting.

### Western blot analysis

Samples containing equal amounts of protein were separated using sodium dodecyl sulfate (SDS)-PAGE and then transferred to polyvinylidene difluoride (PVDF) membranes (Millipore). After blocking with 5% skim milk in TBST for 2 h, membranes were incubated with primary antibodies (diluted 1:1000) for 1 h, followed by incubation with horseradish peroxidase (HRP)-conjugated secondary antibodies (diluted 1:2000; ProteinTech) for 45 min. Membranes were gently soaked in an enhanced chemiluminescence (ECL) solution (Millipore) and protein bands were visualized using a ChemiDoc MP gel imager (Bio-Rad Laboratories, Hercules, CA, USA).

### Membrane fusion assay

293T cells were pre-seeded in 24-well plates pre-coated with poly-D-lysine (Sigma). Seeded cells were co-transfected with pEGFP-N1 and pCAGGS-LASV GPC plasmids using PEI. After 24 h of transfection, cells were incubated in different pH medium for 15 min at 37°C to enable glycoprotein triggering, respectively. The cells were then restored to neutral medium and cultured for an additional 4 h to allow membrane rearrangement. Syncytium formation was manually identified and visualized using a fluorescence microscope (Olympus).

To quantify the efficiency of membrane fusion, a dual-luciferase reporter assay was performed as described previously (25–27). Briefly, 293T cells in a 24-well plate were co-transfected with plasmids expressing T7 RNA polymerase (pCAGT7) and pCAGGS-LASV GPC. 293T cells in a 6-well plate were transfected with pRL-CMV together with pT7EMCVLuc CMV (the three plasmids used in this reporter assay were kindly gifted by Yosh iharu Matsuura, Osaka University, Osaka, Japan). After transfection for 24 h, the cells were gently trypsinized and co-cultured for 6 h in DMEM containing 10% FBS. Membrane fusion was initiated by treatment with acidified medium, which was adjusted using citric acid buffer, and incubated in DMEM supplemented with 2% FBS for 24 h. Membrane fusion activity was quantitatively assessed by detecting the expression of firefly luciferase and sea pansy luciferase using the Dual-Luciferase Reporter Assay System (Promega, Madison, WI, USA) according to the manufacturer’s protocol.

### Production of pseudovirus

Plasmid pVSVΔG-eGFP (Addgene plasmid #31842) was modified to pVSVΔG-Rluc to generate LASV GPC pseudotyped viruses (LASVpv). LASVpv were produced as described previously (25–28). Briefly, 293T cells were transfected with the envelope LASV GPC-pCAGGS plasmid or its mutant derivatives using PEI. After 24 h, the cells were infected with pVSVΔG-Rluc at an MOI of 0.1, for 1 h. Culture supernatants containing viral particles were harvested 30 h later and centrifuged to remove cell debris.

Viral RNA was extracted from 200 μL of pseudovirus using the MiniBEST Viral RNA/DNA Extraction Kit Ver.5.0 (TaKaRa, Japan), and the extracted mRNA was used as a template for reverse transcription using M-MLV reverse transcriptase (Promega, Madison, WI, USA). Virus was quantified using real-time PCR using SYBR Premix Ex Taq™ (Applied Biosystems, Thermo Fisher Scientific, USA) according to the manufacturer’s instructions. Ten-fold serial dilutions of VSV-eGFP were used to construct a standard curve for calculating viral copy numbers.

### Construction of LAMP1-knockout A549 cells

LAMP1-knockout A549 cells were constructed using the CRISPR/Cas9 system. The primers covering the cleavage site on both strands were as follows: forward primer, CACCGAACGGGACCGCGTGCATAA and reserve primer, AAACTTATGCACGCGGTCCCGTTC. A549 cells (5 × 10^5^) were seeded into one well of a six-well plate and were transfected with 600 ng of plasmid with 5 μL of Lipofectamine 2000 (Invitrogen) in 200 μL of OptiMEM (Gibco). At 48 h post transfection, fresh medium containing 2 μg/mL puromycin was added, and the cells were incubated for 2 days. Puromycin-resistant A549 cells were then diluted and cloned. Finally, single-cell clones were identified, amplified, genotyped, and cultured for further experiments. LAMP1 expression in the knockout line was evaluated using western blotting with an anti-LAMP1 antibody.

### Infectivity assay

To measure the effect of LAMP1 knockout on the infectivity of wild-type and mutant LASVpv, BHK, HEK 293T, and Vero cells, as well as wild-type and KO-LAMP1 A549 cells were seeded at a density of 1.5 × 10^5^/mL into 96-well plates and then infected 16 h later with the indicated virus (2 × 10^6^ viral copy number). Supernatants were removed at 24 h p.i., and cell lysates were collected to perform luciferase assays.

### Kinetics assay by BLI

The distal LAMP1-Fc fusion protein was biotinylated in a buffer containing 50 mM sodium citrate (pH 5.0) and 0.02% Tween. The interaction between distal-LAMP1 protein and GP1_LASV_ was assessed using BLI on an Octet RED96 instrument (Pall ForteBio LLC, CA). Biotinylated distal LAMP1 (50 μg/mL) was immobilized on the streptavidin-coated biosensors, then four concentrations (2-fold dilutions; 200-1600 nM) of GP1_LASV_-Fc protein were used for detection. The experiments consisted of five steps: baseline acquisition, distal LAMP1 protein loading onto the sensor, second baseline acquisition, association of GP1_LASV_ protein, and dissociation of GP1_LASV_ protein. The baseline and dissociation steps were assessed using kinetic buffer. The ability of GP1_LASV_ to bind to LAMP1 was determined using response value measurements (12). Experimental data were analyzed using Octet data acquisition software v.7.1, and curve fitting was performed using GraphPad prism (29).

### Statistical analysis

Statistical analyses were performed using GraphPad Prism 7 software (GraphPad Inc., La Jolla, CA, USA). All statistical analyses were performed using Student’s t-test.

## Acknowledgements

We thank Professor Ron Diskin (Department of Structural Biology, Weizmann Institute of Science, Rehovot, Israel) for providing technical details and plasmids. We thank the Center for Instrumental Analysis and Metrology, Core Facility and Technical Support, Wuhan Institute of Virology, for providing technical assistance. This work was supported by the National Key Research and Development Program of China (2018YFA0507204), the National Natural Sciences Foundation of China (31670165), Wuhan National Biosafety Laboratory, Chinese Academy of Sciences Advanced Customer Cultivation Project (2019ACCP-MS03), and the Open Research Fund Program of the State Key Laboratory of Virology of China (2018IOV001).

